# A CRISPR-based strategy for targeted sequencing in biodiversity science

**DOI:** 10.1101/2023.06.30.547247

**Authors:** Bethan Littleford-Colquhoun, Tyler R. Kartzinel

## Abstract

Many applications in molecular ecology require the ability to match specific DNA sequences from single- or mixed-species samples to a diagnostic reference library. Widely used methods for DNA barcoding and metabarcoding require PCR and amplicon sequencing to identify taxa based on target sequences, but the target-specific enrichment capabilities of CRISPR-Cas systems may offer advantages in some applications. We identified 54,837 CRISPR-Cas guide RNAs that may be useful for enriching chloroplast DNA across phylogenetically diverse plant species. We then tested a subset of 17 guide RNAs *in vitro* to enrich and sequence plant DNA strands ranging in size from diagnostic DNA barcodes of 1,428 bp to entire chloroplast genomes of 121,284 bp. We used an Oxford Nanopore sequencer to evaluate sequencing success based on both single- and mixed-species samples, which yielded mean on-target chloroplast sequence lengths of 5,755-11,367 bp, depending on the experiment. Single-species experiments yielded more on-target sequence reads and greater accuracy, but mixed-species experiments yielded superior coverage. Comparing CRISPR-based strategies to a widely used protocol for plant DNA metabarcoding with the chloroplast *trn*L-P6 marker, we obtained a 66-fold increase in sequence length and markedly better estimates of relative abundance for a commercially prepared mixture of plant species. Future work would benefit from developing both *in vitro* and *in silico* methods for analyses of mixed-species samples, especially when the appropriate reference genomes for contig assembly cannot be known *a priori*. Prior work developed CRISPR-based enrichment protocols for long-read sequencing and our experiments pioneered its use for plant DNA barcoding and chromosome assemblies that may have advantages over workflows that require PCR and short-read sequencing.

## Introduction

In biomedical science, CRISPR-Cas systems are regularly used to target a section of DNA with high precision and accuracy (Kaminski et al., 2021; Wang et al., 2022). Although most applications of CRISPR have utilized its genome-editing capabilities, its target-specific binding and cutting capabilities for DNA detection and enrichment are increasingly evident (Phelps et al., 2020). To date, CRISPR has been used: to detect the presence of specific genes of interest such as antibiotic resistance in *Staphylococcus* (Quan et al., 2019), drug resistance in the malaria parasite *Plasmodium falciparum* (Cunningham et al., 2021), and SARS-Cov-2 (Broughton et al., 2020); to identifying SNPs associated with lung cancer (Qiu et al., 2018) and the hepatitis B virus (Ke et al., 2021), and bacterial genes within environmental samples (Sandoval-Quintana et al., 2023). While CRISPR-Cas systems are becoming the most reliable, affordable, and versatile method for analyzing DNA, they may be generally underutilized in environmental biology (Phelps et al., 2020).

What makes CRISPR so reliable and versatile is its ability to recognize a target sequence with high precision (Knott & Doudna, 2018). Type II CRISPR-Cas systems are the best characterized and most commonly used (Xu & Li, 2020), comprising two key components: a guide RNA (gRNA), which recognizes the target sequence, and a CRISPR-associated endonuclease (Cas protein) that cuts the targeted sequence. Guide RNAs are composed of ‘scaffold sequences’ necessary for Cas-binding and user-defined ∼20 nucleotide spacer sequences that correspond to a ‘target sequence’ to be cleaved from template DNA by the Cas-gRNA ribonucleoprotein (RNP) complex. Much like designing PCR primers, specific sequences can be targeted based on the spacer sequence of the gRNA. Similarly, the gRNA can tolerate some degree of mismatching between target and spacer sequences: the gRNA binds to the target in a 3′ to 5′ direction such that mismatches at the 3′ end of the spacer can prevent cleavage whereas ∼2 bp mismatches toward the 5′ end may often be tolerated (Fu et al., 2016). Unlike PCR primers, however, the target sequence must be located immediately before nuclease-specific protospacer adjacent motif (PAM) sequences that are numerous throughout the genome and required for Cas nuclease cleavage. When gRNAs can be synthesized for a target locus, the result is not many copies of the target (as in PCR) but rather enrichment of the target as it is cleaved directly from the template.

A distinct benefit to using CRISPR-Cas enrichment is that, unlike PCR, many (100+) gRNAs can be multiplexed within a single assay (Gilpatrick et al., 2023; López-Girona et al., 2020; Xie et al., 2015), allowing for multiple regions within a genome to be enriched in a single reaction. Multiple scoring models have been developed to help identify gRNAs with high efficiency (on-target activity) and specificity (limited off-target activity), though they vary in their intended uses and thus it has been historically challenging to translate their utility beyond model systems (Cui et al., 2018; Sledzinski et al., 2020; Wilson et al., 2018). As the ability to design effective gRNAs continues to improve, many more of these potential applications in biodiversity research may begin to be realized (Gilpatrick et al., 2023).

Both DNA barcoding and metabarcoding are widely used strategies that rely on PCR to enrich sequences from single- or mixed-species samples, respectively, in order to compare the resulting sequences with reference data (Srivathsan et al., 2021). Unfortunately, existing DNA reference databases are often biased towards certain markers and it has been difficult to achieve consensus about which barcodes to use for certain taxa, in part because reliance on PCR limits the length of target sequences in ways that can constrain taxonomic precision (CBOL Plant Working Group, 2009; Deck et al., 2012; Hebert et al., 2022; Hoban et al., 2022; Keck et al., 2022). When similar approaches are applied to samples containing mixtures of DNA from multiple species using DNA metabarcoding, reliance on PCR involves further challenges associated with detecting and estimating the relative abundance of phylogenetically disparate taxa (Clarke et al., 2014; Deagle et al., 2019; Kelly et al., 2019; O’Donnell et al., 2016; Stapleton et al., 2022). By contrast, CRISPR-Cas technology may enable researchers to circumvent several of these challenges: it has recently enabled the targeted enrichment of entire mitogenomes from diverse fishes (Ramon-Laca et al., 2022); it has enabled the detection of specific DNA strands in environmental DNA (Baerwald et al., 2023; Karlikow et al., 2023; Sánchez et al., 2022; Williams et al., 2023; Williams et al., 2021; Williams et al., 2019); and it has revealed both single-nucleotide and structural variants in the locus controlling for fruit color in apples (López-Girona et al., 2020). Yet despite the potential to use CRISPR-Cas to overcome drawbacks to PCR by providing longer, and hence more diagnostic markers, this strategy has not yet been tested in comparative analyses involving multiple target sequences.

We developed a set of novel CRISPR-based protocols to enrich plant DNA barcodes. We began by evaluating the availability of gRNA sequences capable of targeting chloroplast DNA across a broad swath of the angiosperm phylogeny, which should enable ‘universal’ DNA enrichment strategies. Then we designed protocols to target the enrichment of CRISPR-associated loci *in vitro*. We evaluated the strengths and weaknesses of broad-spectrum strategies for enriching markers that ranged in size from 1,428 bp to the entire chloroplast genome from single- or mixed-species samples by: (*i*) comparing three strategies for enriching standard plant DNA barcode loci from a single species DNA sample, (*ii*) enriching a whole chloroplast to assemble a reference genome for a single species, (*iii*) applying a barcode-enrichment strategy to a mixed sample of known species composition.

## Methods

### Assessing cross-species coverage of guide RNAs (gRNAs)

Our goal was to identify broad-spectrum gRNAs that targeted chloroplast DNA sequences from many species. We designed gRNAs for the Type II CRISPR-Cas system. This system relies on the Cas9 (SpCas9) protein which recognizes a PAM sequence of NGG in a 5′ to 3′ direction (where ‘N’ can be any nucleotide base). We began by identifying candidate gRNAs that appeared in chloroplast reference genomes across a set of 7 well-studied, economically important, and phylogenetically disparate plant species: three grasses (wheat, *Triticum aestivum*; oats, *Avena sativa*; corn, *Zea mays*), two superrosids (soybeans, *Glycine max*; peanuts, *Arachis hypogaea*), and two superasterids (sunflower, *Helianthus annuus*; spinach, *Spinacia oleracea*; see Table 2 for RefSeq accession numbers). We did this by searching for all potential gRNAs in the chloroplast reference genomes of the 7 target species using the *Find CRISPR site* tool within Geneious Prime 2023.0.4. We evaluated the predicted *in vitro* functionality of these gRNAs based on features including GC count, position-independent nucleotide counts, the location of the gRNA target site within the gene, and the thermodynamic properties of each identified gRNA using *Rule Set 2* (Doench et al., 2016). The *Rule Set 2* model gives high scores to candidate gRNAs that are predicted to efficiently guide Cas9 to the correct spot for cleavage (i.e., on-target activity), enabling comparisons of the candidate gRNAs across genomic sites and target taxa. Once candidate gRNAs were identified using each reference genome independently, we tallied the number of references that contained an exact (100%) match between guide and the target sequences (i.e., assuming strict, no tolerance for mismatches). We evaluated (*i*) how many exact gRNAs were present in only one species (i.e., narrowest coverage), (*ii*) how many unique gRNAs occurred across all species (i.e., broadest coverage), and (*iii*) how many gRNAs had multiple match sites within a species (i.e., poor site fidelity). Then we identified gRNAs that exactly matched ≥5 of the 7 phylogenetically disparate target species, tolerating up to 2 mismatches at the 5′ end of the gRNA. Finally, we selected candidate gRNAs that had good predicted *in vitro* functionality and coverage across the 7 target species based on the criteria outlined above and had a *Rule Set 2* score ≥0.2 in all target species.

### Selection of gRNAs and in vitro testing

We selected a subset of broad-coverage candidate gRNAs for use in a series of six experiments to: (*i*) sequence standard plant DNA barcode loci and (*ii*) a complete chloroplast genome from a DNA sample representing a single species (spinach) as well as to (*iii*) elucidate the sequence composition of a mixed-sample containing six known plant species (wheat, oats, corn, soybean, peanuts, sunflower; Figure 1). We refer to these overarching strategies, intuitively, as the ‘barcoding approach’, ‘whole chloroplast approach’, and ‘mixed-species approach’.

**Figure 1.**
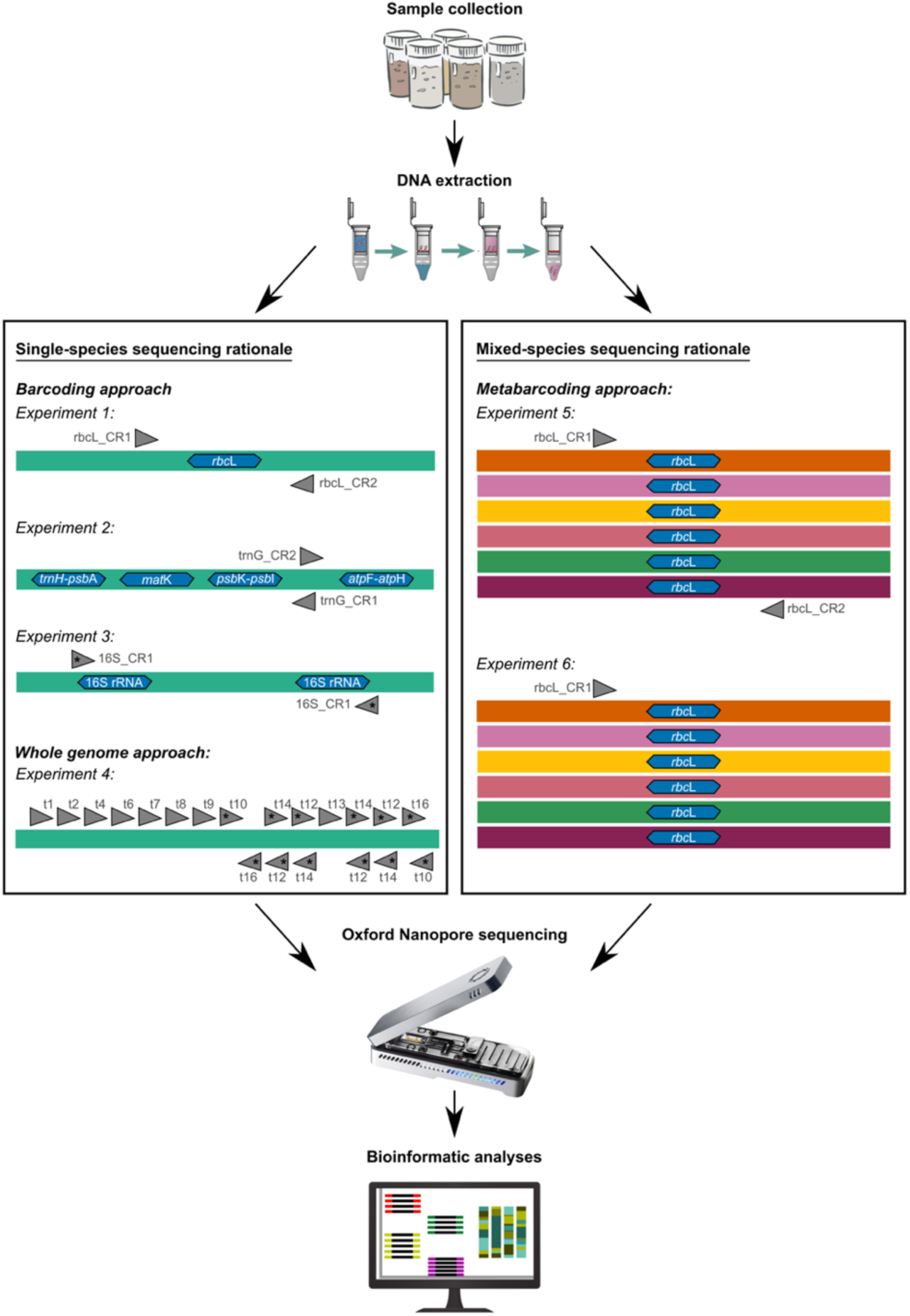
Experimental overview for single- and mixed-species sequencing approaches. All experiments began with sample collection and DNA extraction (top) and ended with Oxford Nanopore MinION sequencing followed by bioinformatic analyses (bottom). Experiments differed according to the strategy for designing gRNAs used for enrichment (box). The colored bands represent DNA strands from each of the seven species used across these experiments, the blue hexagons identify the barcode markers targeted in each experiment, and gray arrowheads show the location and directionality of each gRNA binding site (Table 1). Asterisks at the biding sites indicate gRNAs that occur at ≥2 locations within the chloroplast, as expected for the inverted repeats of 16S rRNA.

**Table 1.**
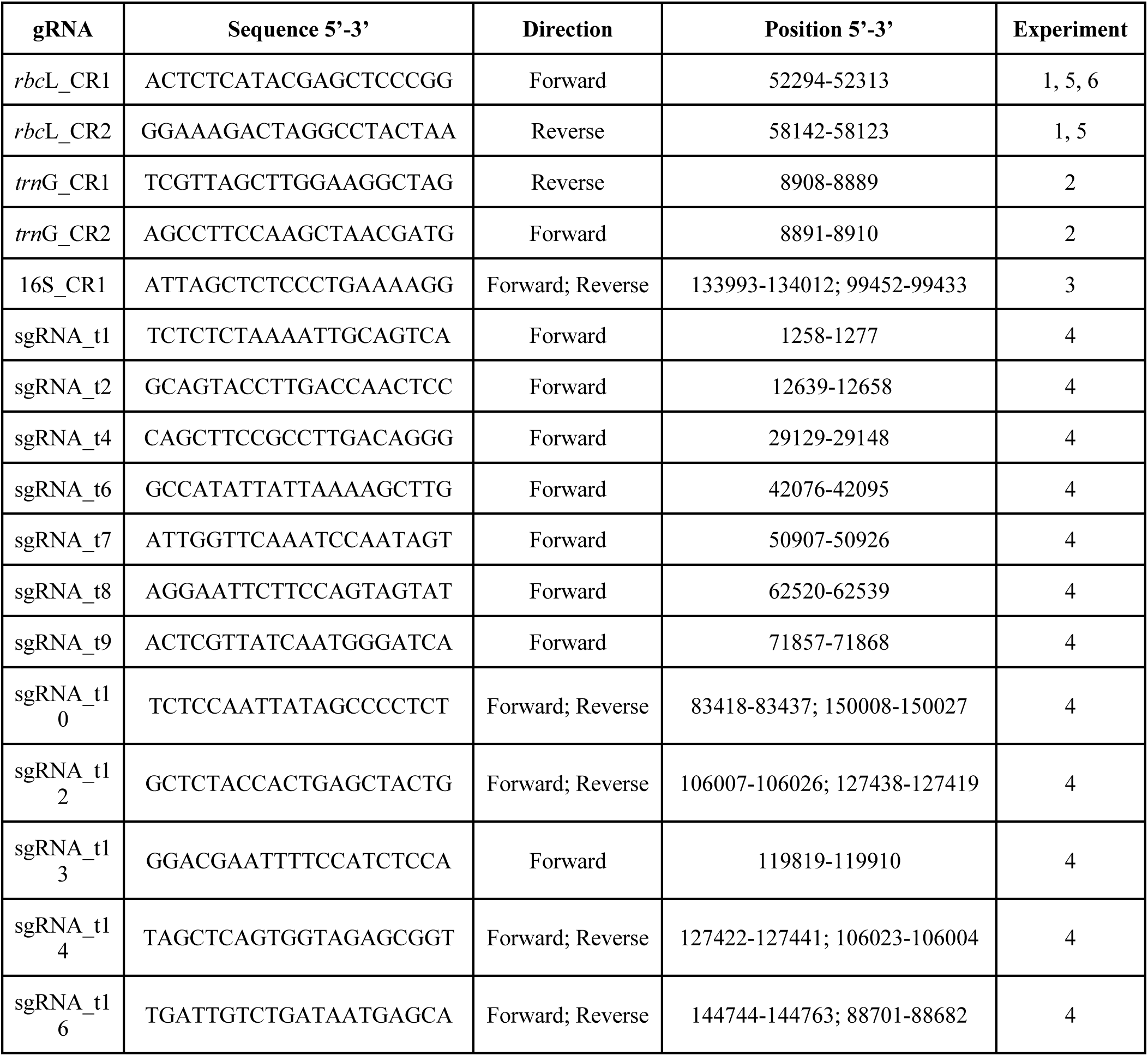
Guide RNA (gRNA) sequences used to direct CRISPR-Cas9 scission in Experiments 1-6. For each gRNA, we provide a unique identifier, the sequence, the direction of activity (“forward” direction indicates that the protospacer adjacent motif (PAM) and target sequence is found upstream of the region of interest on the forward DNA strand; “reverse” indicates that the PAM and target sequence is found downstream of the region of interest on the reverse DNA strand), the position of the gRNA with respect to the spinach reference chloroplast genome (Table 2), and the experiment(s) for which we trialed each gRNA (Figure 1).

**Table 2.**
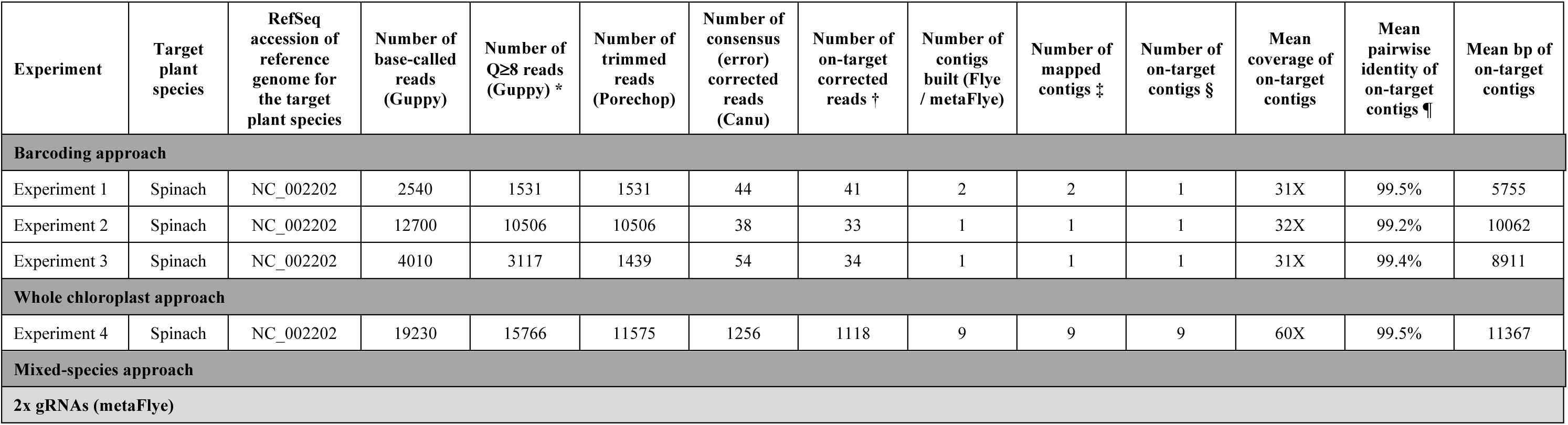
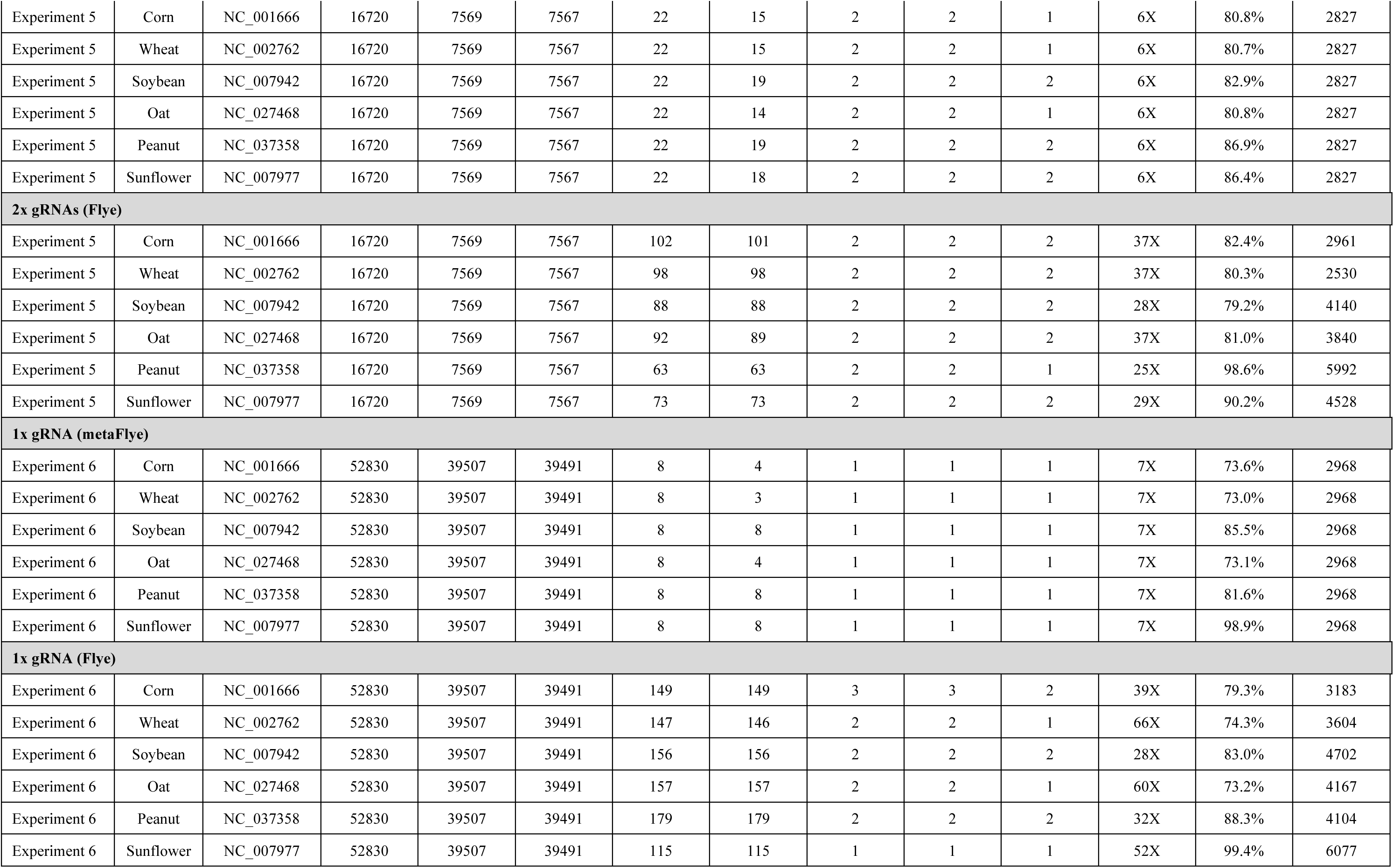
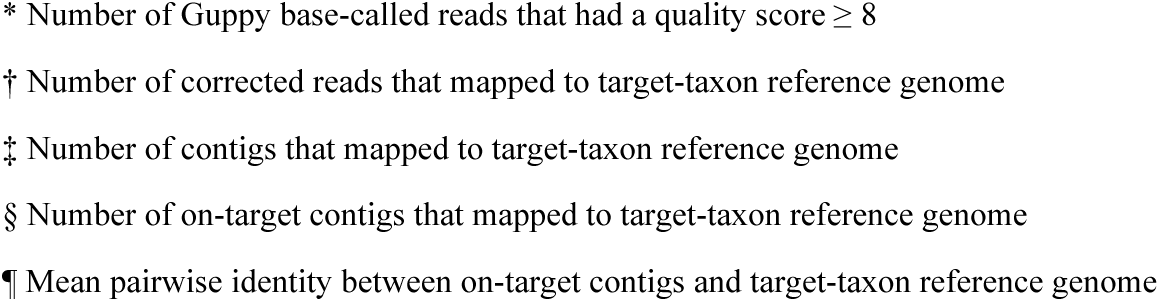
Results from each step in bioinformatic pipeline. For experiments 1-6, we provide the target plant species and the accession number of the corresponding reference genome used to align output reads, the total number of Guppy base-called reads, the number of reads that passed Guppy quality control, the number of reads retained following adapter trimming in Porechop, the number of consensus (error) corrected reads produced using Canu, the number of corrected reads that mapped to the region of interest (on-target), the number of contigs assembled from these corrected reads using Flye/metaFlye, the number of assembled contigs that mapped to the target species reference genome, the number of assembled contigs mapped to the region of interest (on-target), and the mean fold-coverage, mean pairwise identity, and the length of all on-target contigs. Outputs for different approaches (barcoding, whole chloroplast, and mixed-species) are shown using dark grey banners. For the mixed-species approach (Experiments 5-6), we report separate outputs (light grey banners) depending on whether contigs were assembled using all corrected reads of a sequencing run (using metaFlye) or whether contigs were assembled independently for each target taxon (using Flye).

| Experiment | Target plant species | RefSeq accession of reference genome for the target plant species | Number of base-called reads (Guppy) | Number of Q≥8 reads (Guppy) * | Number of trimmed reads (Porechop) | Number of consensus (error) corrected reads (Canu) | Number of on-target corrected reads † | Number of contigs built (Flye / metaFlye) | Number of mapped contigs ‡ | Number of on-target contigs § | Mean coverage of on-target contigs | Mean pairwise identity of on-target contigs ¶ | Mean bp of on-target contigs |
| --- | --- | --- | --- | --- | --- | --- | --- | --- | --- | --- | --- | --- | --- |
| **Barcoding approach** |  |  |  |  |  |  |  |  |  |  |  |  |  |
| Experiment 1 | Spinach | NC_002202 | 2540 | 1531 | 1531 | 44 | 41 | 2 | 2 | 1 | 31X | 99.5% | 5755 |
| Experiment 2 | Spinach | NC_002202 | 12700 | 10506 | 10506 | 38 | 33 | 1 | 1 | 1 | 32X | 99.2% | 10062 |
| Experiment 3 | Spinach | NC_002202 | 4010 | 3117 | 1439 | 54 | 34 | 1 | 1 | 1 | 31X | 99.4% | 8911 |
| **Whole chloroplast approach** |  |  |  |  |  |  |  |  |  |  |  |  |  |
| Experiment 4 | Spinach | NC_002202 | 19230 | 15766 | 11575 | 1256 | 1118 | 9 | 9 | 9 | 60X | 99.5% | 11367 |
| **Mixed-species approach** |  |  |  |  |  |  |  |  |  |  |  |  |  |
| **2x gRNAs (metaFlye)** |  |  |  |  |  |  |  |  |  |  |  |  |  |
| Experiment 5 | Corn | NC_001666 | 16720 | 7569 | 7567 | 22 | 15 | 2 | 2 | 1 | 6X | 80.8% | 2827 |
| Experiment 5 | Wheat | NC_002762 | 16720 | 7569 | 7567 | 22 | 15 | 2 | 2 | 1 | 6X | 80.7% | 2827 |
| Experiment 5 | Soybean | NC_007942 | 16720 | 7569 | 7567 | 22 | 19 | 2 | 2 | 2 | 6X | 82.9% | 2827 |
| Experiment 5 | Oat | NC_027468 | 16720 | 7569 | 7567 | 22 | 14 | 2 | 2 | 1 | 6X | 80.8% | 2827 |
| Experiment 5 | Peanut | NC_037358 | 16720 | 7569 | 7567 | 22 | 19 | 2 | 2 | 2 | 6X | 86.9% | 2827 |
| Experiment 5 | Sunflower | NC_007977 | 16720 | 7569 | 7567 | 22 | 18 | 2 | 2 | 2 | 6X | 86.4% | 2827 |
| **2x gRNAs (Flye)** |  |  |  |  |  |  |  |  |  |  |  |  |  |
| Experiment 5 | Corn | NC_001666 | 16720 | 7569 | 7567 | 102 | 101 | 2 | 2 | 2 | 37X | 82.4% | 2961 |
| Experiment 5 | Wheat | NC_002762 | 16720 | 7569 | 7567 | 98 | 98 | 2 | 2 | 2 | 37X | 80.3% | 2530 |
| Experiment 5 | Soybean | NC_007942 | 16720 | 7569 | 7567 | 88 | 88 | 2 | 2 | 2 | 28X | 79.2% | 4140 |
| Experiment 5 | Oat | NC_027468 | 16720 | 7569 | 7567 | 92 | 89 | 2 | 2 | 2 | 37X | 81.0% | 3840 |
| Experiment 5 | Peanut | NC_037358 | 16720 | 7569 | 7567 | 63 | 63 | 2 | 2 | 1 | 25X | 98.6% | 5992 |
| Experiment 5 | Sunflower | NC_007977 | 16720 | 7569 | 7567 | 73 | 73 | 2 | 2 | 2 | 29X | 90.2% | 4528 |
| **1x gRNA (metaFlye)** |  |  |  |  |  |  |  |  |  |  |  |  |  |
| Experiment 6 | Corn | NC_001666 | 52830 | 39507 | 39491 | 8 | 4 | 1 | 1 | 1 | 7X | 73.6% | 2968 |
| Experiment 6 | Wheat | NC_002762 | 52830 | 39507 | 39491 | 8 | 3 | 1 | 1 | 1 | 7X | 73.0% | 2968 |
| Experiment 6 | Soybean | NC_007942 | 52830 | 39507 | 39491 | 8 | 8 | 1 | 1 | 1 | 7X | 85.5% | 2968 |
| Experiment 6 | Oat | NC_027468 | 52830 | 39507 | 39491 | 8 | 4 | 1 | 1 | 1 | 7X | 73.1% | 2968 |
| Experiment 6 | Peanut | NC_037358 | 52830 | 39507 | 39491 | 8 | 8 | 1 | 1 | 1 | 7X | 81.6% | 2968 |
| Experiment 6 | Sunflower | NC_007977 | 52830 | 39507 | 39491 | 8 | 8 | 1 | 1 | 1 | 7X | 98.9% | 2968 |
| **1x gRNA (Flye)** |  |  |  |  |  |  |  |  |  |  |  |  |  |
| Experiment 6 | Corn | NC_001666 | 52830 | 39507 | 39491 | 149 | 149 | 3 | 3 | 2 | 39X | 79.3% | 3183 |
| Experiment 6 | Wheat | NC_002762 | 52830 | 39507 | 39491 | 147 | 146 | 2 | 2 | 1 | 66X | 74.3% | 3604 |
| Experiment 6 | Soybean | NC_007942 | 52830 | 39507 | 39491 | 156 | 156 | 2 | 2 | 2 | 28X | 83.0% | 4702 |
| Experiment 6 | Oat | NC_027468 | 52830 | 39507 | 39491 | 157 | 157 | 2 | 2 | 1 | 60X | 73.2% | 4167 |
| Experiment 6 | Peanut | NC_037358 | 52830 | 39507 | 39491 | 179 | 179 | 2 | 2 | 2 | 32X | 88.3% | 4104 |
| Experiment 6 | Sunflower | NC_007977 | 52830 | 39507 | 39491 | 115 | 115 | 1 | 1 | 1 | 52X | 99.4% | 6077 |

\* Number of Guppy base-called reads that had a quality score $\geq 8$
† Number of corrected reads that mapped to target-taxon reference genome
‡ Number of contigs that mapped to target-taxon reference genome
§ Number of on-target contigs that mapped to target-taxon reference genome
¶ Mean pairwise identity between on-target contigs and target-taxon reference genome

We began with the relatively simple, single species ‘barcoding approach’ using spinach as the target species (Experiments 1-3; Figure 1). We targeted the three ‘standard’ plant DNA barcodes as well as other markers that have been considered potentially useful for DNA barcoding (CBOL Plant Working Group, 2009; Kress, 2017). Experiment 1 used 2 gRNAs to target the whole *rbc*L gene (1,428 bp), which includes the standard *rbc*L barcode locus (553 bp; CBOL Plant Working Group, 2009; Kress, 2017). One gRNA targeted a region upstream of the barcode (the ‘forward’ gRNA) and one targeted a region downstream of the barcode (‘reverse’) such that the two gRNA binding sites were separated by 5,809 bp (Figure 1, Table 1). Experiment 2 targeted multiple DNA barcodes by making a two-directional break in the *trn*G gene with forward and reverse gRNAs that overlapped by 18 bp (Figure 1, Table 1). Within spinach, four potentially useful plant DNA barcodes sit within 9,000 bp of *trn*G and can be targeted for sequencing in this way (the standard *mat*K and *trn*H-*psb*A barcodes as well as *psb*K-*psb*I, and *atp*F-*atp*H; CBOL Plant Working Group, 2009; Kress, 2017). Experiment 3 targeted the inverted 16S rRNA repeat region (1,491 bp) of the chloroplast genome and aimed to determine whether we could use a single gRNA to sequence both regions, because 16S is a structurally interesting region of the chloroplast (Manhart, 1995; Strauss et al., 1988) even though it is more often targeted as a DNA barcode for other taxa [e.g., bacteria (Caporaso et al., 2012), animals (Kartzinel & Pringle, 2015; Vences et al., 2005); Figure 1, Table 1].

Our second overarching aim was to test methods suitable for a ‘whole chloroplast approach’ using spinach as the target species (Experiment 4, Figure 1). We used 12 gRNAs that were predicted to enable enrichment around 20 cut sites that were relatively evenly spaced throughout the spinach chloroplast (∼5,200-17,500 bp apart). Of the 12 gRNAs that we selected, 8 occurred only once in the spinach reference chloroplast (forward in directionality), 2 occurred twice (only when mismatches were tolerated), and 2 occurred 4 times as they were located in the large inverted rRNA subunit repeats (Figure 1, Table 1).

Finally, we selected candidate gRNAs to test a ‘mixed-species approach’ for identifying taxa (Experiments 5-6; Figure 1). These experiments aimed to sequence 6 target species that were ground and pelleted by the commercial supplier Teklad Lab Animal Diets. These pellets represented a homogeneous mixture comprising Teklad Global Rodent 2016 formula (wheat, corn, soybean; TD.00217) and 3 additional plant components (oats, peanuts, sunflowers) that were mixed in even proportion. The overall ratios were 50% Teklad Global Rodent 2016 to 50% additional plant components, resulting in hypothetical plant biomass ratios of oat, peanut, and sunflower at 1/6^th^ each plus wheat, corn, and soy that each comprised an unspecified (proprietary) ratio within the other 50%; because the core Teklad components were processed more intensively than the additional components we assumed a greater level of DNA degradation in the former than the latter. Experiments 5 and 6 both targeted *rbc*L, but Experiment 5 used a pair of forward and reverse gRNAs while Experiment 6 used only a forward gRNA (Figure 1, Table 1). Due to chloroplast rearrangements, the gRNAs appear at different genomic locations across the six reference chloroplast genomes but are linked to the *rbc*L locus in each. We compared Experiments 5 and 6 to determine which provided a greater number of on-target sequences and a better estimate of plant DNA relative read abundance (RRA) in the mixture.

### Sequencing library preparation

Total genomic DNA was extracted from ∼0.2 mg (*i*) spinach and (*ii*) mixed-species Teklad samples using a Zymo Quick-DNA Fecal/Soil Microbe Miniprep Kit (Zymo Research). Following extractions, we quantified DNA using a Qubit dsDNA high sensitivity assay kit (Invitrogen). For each experiment, we enriched the target chloroplast regions using the nanopore Cas9-targeted sequencing method (nCATS; Gilpatrick et al., 2020). Briefly, this method uses Cas9-mediated DNA cleavage to cut double-stranded DNA ∼3-4 nucleotides upstream of the target PAM sequence. This enables us to enrich target DNA by selectively ligating adapters to the cut sites created by the Cas9/gRNA–ribonucleoprotein (RNP) complexes created in each experiment. We built custom gRNA duplexes for each experiment and assembled them into the RNP complex by adding 1 μl of pooled crRNAs (user-defined spacer sequences; IDT) and 1 μl of Alt-R^®^ CRISPR-Cas9 tracrRNA (IDT) to 8 μl of nuclease free water and incubating at 95°C for 5 minutes. The RNP complex was then created by incubating 1.2 μl Alt-R^®^ HiFi Cas9 Nuclease (IDT), 2.8 μl 10X CutSmart Buffer (NEB), 23 μl nuclease free water and 3 μl of the gRNA duplex at room temperature for 20 minutes. To selectively enrich the region of interest, we first dephosphorylated pre-existing DNA ends before cutting with Cas9 to preferentially ligate sequencing adapters to the cut sites created by the RNP complex. We did this by incubating 1.5 ng of genomic DNA, 3 μl 10X CutSmart buffer, and 3 μl QuickCIP enzyme (NEB) at 37°C for 10 minutes, followed by enzyme inactivation at 80°C for 2 minutes. Cleavage and dA-tailing of the dephosphorylated DNA occurred in a reaction using 10 μl of the assembled RNP complex, 10 mM dATP (Zymo Research), and 1 μl Taq DNA polymerase (NEB) with incubation at 37°C for 15 minutes followed by 72°C for 5 minutes.

### Oxford Nanopore sequencing

For each experiment, we sequenced the enriched target loci used long-read nanopore sequencing (Gilpatrick et al., 2020). Nanopore sequencing adapters were first ligated to Cas9 cut sites by incubating 10 μl of NEBNext Quick T4 DNA Ligase (NEB), 20 μl ligation buffer (ONT), 4.5 μl nuclease free water, and 3.5 μl AMX sequencing adapters (LSK109 sequencing kit; ONT) at room temperature for 10 minutes. An equivolume amount of TE buffer was then added to the ligated sample, followed by 0.3X volume addition of AMPure XP beads (Beckman Coulter). The sample was incubated at room temperature for 5 minutes. The supernatant was removed by pipette after placing the sample on a magnetic rack and then the remaining library was purified twice using 200 μl long-fragment buffer (ONT). We eluted the ligated sample by adding 15 μl elution buffer and incubating at room temperature for 30 minutes before separating the eluate from beads on the magnetic rack. To ensure the recommended 5-50 fmol of library DNA was available for sequencing, we checked library concentrations with a Qubit dsDNA high sensitivity kit. The resulting libraries were sequenced on a MinION Mk1B nanopore sequencer (ONT) using FLO-MIN106D (R9.4.1) flow cells. We added 37.5 μl sequencing buffer (ONT) and 25.5 μl loading beads to the eluate, and then prepared the flow cell by placing 30 μl flush tether (ONT) into a tube of flush buffer (ONT), pulling 230 μl buffer from the priming port, and loading an initial 800 μl of the priming mix. After 5 minutes, an additional 200 μl of priming mix was loaded before the DNA library. The DNA library was added via the SpotON sample port in a dropwise fashion. Finally, we initiated sequencing runs using MinKNOW software (version 22.08.9; ONT), enabling raw data to be processed with *fast basecalling* using Guppy 6.2.11.

### Oxford Nanopore read assembly

First, adapters were trimmed from all reads that passed the Guppy basecaller quality score (Q ≥ 8) using Porechop (Wick et al., 2017). To obtain consensus sequences from overlapping reads, trimmed reads were corrected using the *correct* parameters in Canu with default nanopore settings (Koren et al., 2017). Canu requires information on the expected genome size so that coverage of the input reads can be determined; we used the size of the spinach chloroplast reference genome as the expected size in the whole genome approach (Experiment 4) and the expected target sequence length between gRNAs in all other experiments. The resulting sequences were then assembled *de novo* using Flye v2.9 (Kolmogorov et al., 2019). For mixed-species samples (Experiments 5-6), we used the metagenome assembly mode in Flye (metaFlye) with the *meta* and the *nano-corr* parameters as appropriate for error-corrected nanopore reads. To determine the percent identity between each contig and the reference genome, we mapped contigs to the reference genome(s) of target-species(s) using minimap2 (Li, 2018) with the Oxford Nanopore option in Geneious Prime. When mapping contigs from the 16S rRNA (Experiment 3) and whole genome (Experiment 4) experiments to the spinach chloroplast, we enabled secondary alignments to allow reads to be mapped to multiple locations within the inverted repeats.

### Comparison of CRISPR-Cas enrichment and PCR-based DNA metabarcoding

We compared the hypothetical DNA sequence relative read abundance of target plant species in the mixed-species sample with empirical data obtained using both CRISPR-Cas and PCR-based sequencing approaches (Appendix S1). For the PCR-based benchmark, we used 2 x 150 bp Illumina sequencing and required a strict 100% identity between the resulting amplicon sequences and a global reference library (Appendix S1). To calculate relative read abundance using sequences obtained with CRISPR-Cas enrichment, we used the read-count coverage of the contig built using Flye that mapped to the correct location in the reference genome of each target taxon. To calculate relative read abundance from DNA metabarcoding for the mixed-species sample, we converted sequence counts into proportional data. The relative read abundance values resulting from both methods can be interpreted as estimates of the proportional representation of DNA from the target taxa in the sample after accounting for all sources of bias and error, including variation in the tissue content of DNA per unit biomass, tissue homogenization and extraction, amplification and enrichment, sequencing accuracy, and bioinformatic processes.

## Results

### Coverage of guide RNAs (gRNAs)

In total, we identified 54,837 unique gRNA sequences across the reference chloroplast genomes of the 7 target species (9,852-10,817 unique gRNAs per species; Supplementary Data Table 1). This included (*i*) 44,510 ‘narrow-coverage’ gRNAs that were present in only one target species, (*ii*) 398 ‘broad-coverage’ gRNAs that occurred across all 7 target species, (*iii*) 44,647 ‘high-fidelity’ gRNAs that had only a single cut site in ≥1 target species, and (*iv*) 10,190 ‘low-fidelity’ gRNAs that matched multiple sites in ≥1 target species. Of the 44,647 high-fidelity gRNAs, 6,581 perfectly matched (100% identity) the reference chloroplast of at least 2 target species, and 45 of these occurred in all 7 target species (Supplementary Data Table 1). Of 10,190 low-fidelity gRNAs, 3,746 perfectly matched the reference for at least 2 target species and 353 matched all 7 species (Supplementary Data Table 1). Due to structural differences in the chloroplast genomes of Poaceae, many gRNAs that had perfect homology and a single cut site in the reference genomes of wheat, oat, and corn did not appear in spinach, sunflower, soy, or peanut. Nevertheless, we identified many potentially broad-coverage gRNAs, especially when allowing for ≤2 bp mismatches at the 5’ end of the gRNAs. Moreover, 14 of these broad-coverage gRNAs were located within 3 kb of *rbc*L, 16 were located within 3 kb of *mat*K, and 2 were located within 3 kb *trn*L-P6 across all 7 target species. These CRISPR-associated loci present opportunities to develop broad-spectrum enrichment protocols for DNA barcoding and metabarcoding studies.

### Barcoding approach

We obtained high coverage and accuracy sequencing multiple plant DNA barcodes (Experiments 1-3). Experiment 1 targeted *rbc*L using one forward and one reverse gRNA and yielded a total of 1,531 reads with a Q-score ≥8 (Table 2, Figures 2-3). Raw sequence lengths of 130-15,971 bp (mean: 4,004 bp) encompassed the target length of 5,809 bp. Reads were then error-corrected using Canu and 44 consensus reads were produced; a subset of 41 (93%) reads mapped to the reference spinach chloroplast and 35 mapped to *rbc*L (85% of mapped corrected reads; Table 2). *De novo* assembly using Flye generated 2 contigs with lengths of 7,048 and 5,755 bp that both mapped to the spinach reference, but only the shorter contig aligned to the target region (31X coverage with 99.5% pairwise identity; Table 2). The longer contig did not align to the target region but to an upstream region of the chloroplast (8X coverage).

**Figure 2.**
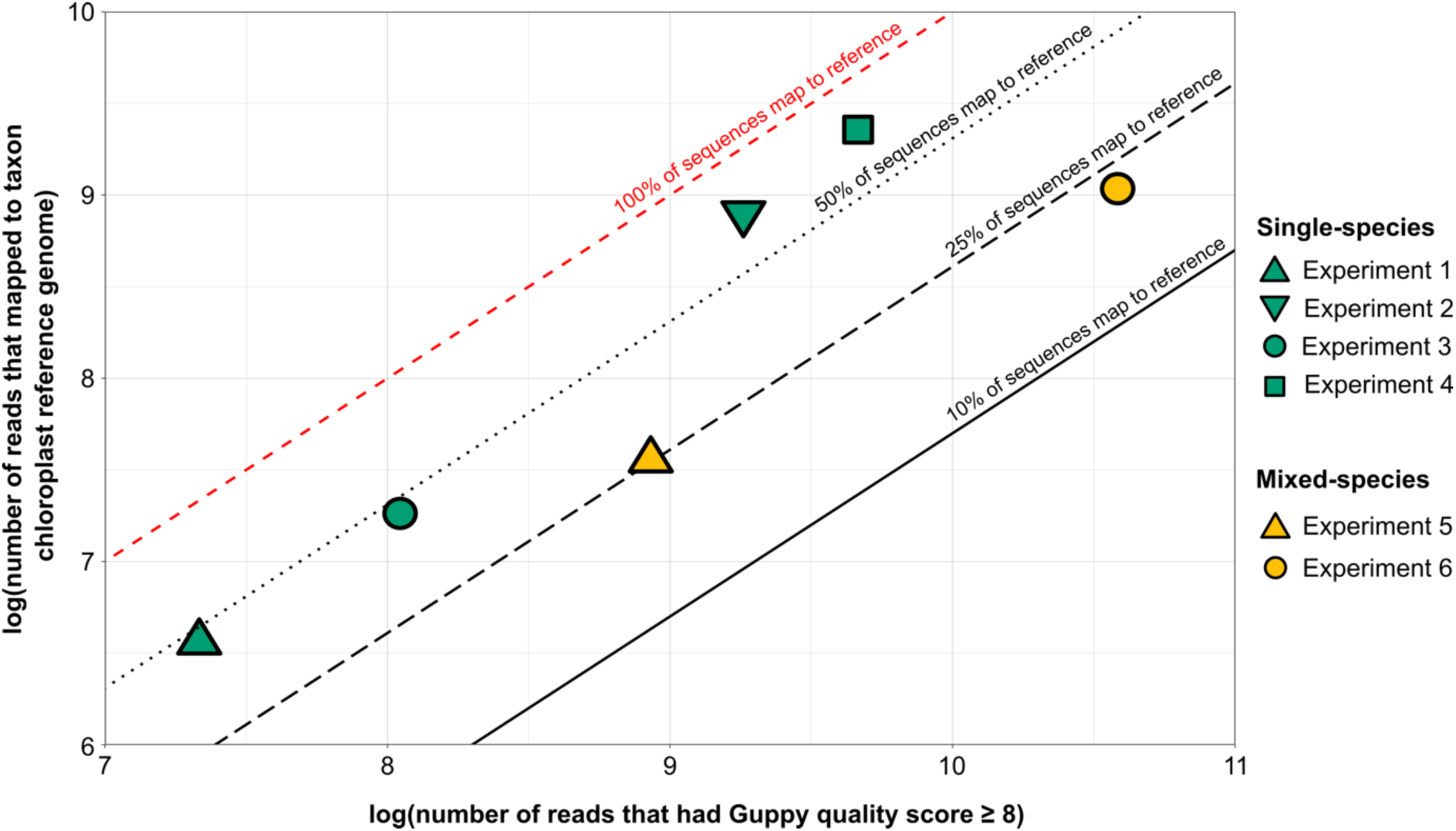
Comparison of the sequence reads generated and successfully mapped to reference genomes in each experiment. The number of DNA sequence reads that passed the Guppy basecaller (Q ≥8) is shown on the x-axis and the number of those reads that mapped to the appropriate chloroplast reference genomes is shown on the y-axis. The four single-species experiments had a greater proportion of base-called reads that mapped to the reference chloroplast genome of the target-taxon compared to mixed-species experiments which included six target taxa.

**Figure 3.**
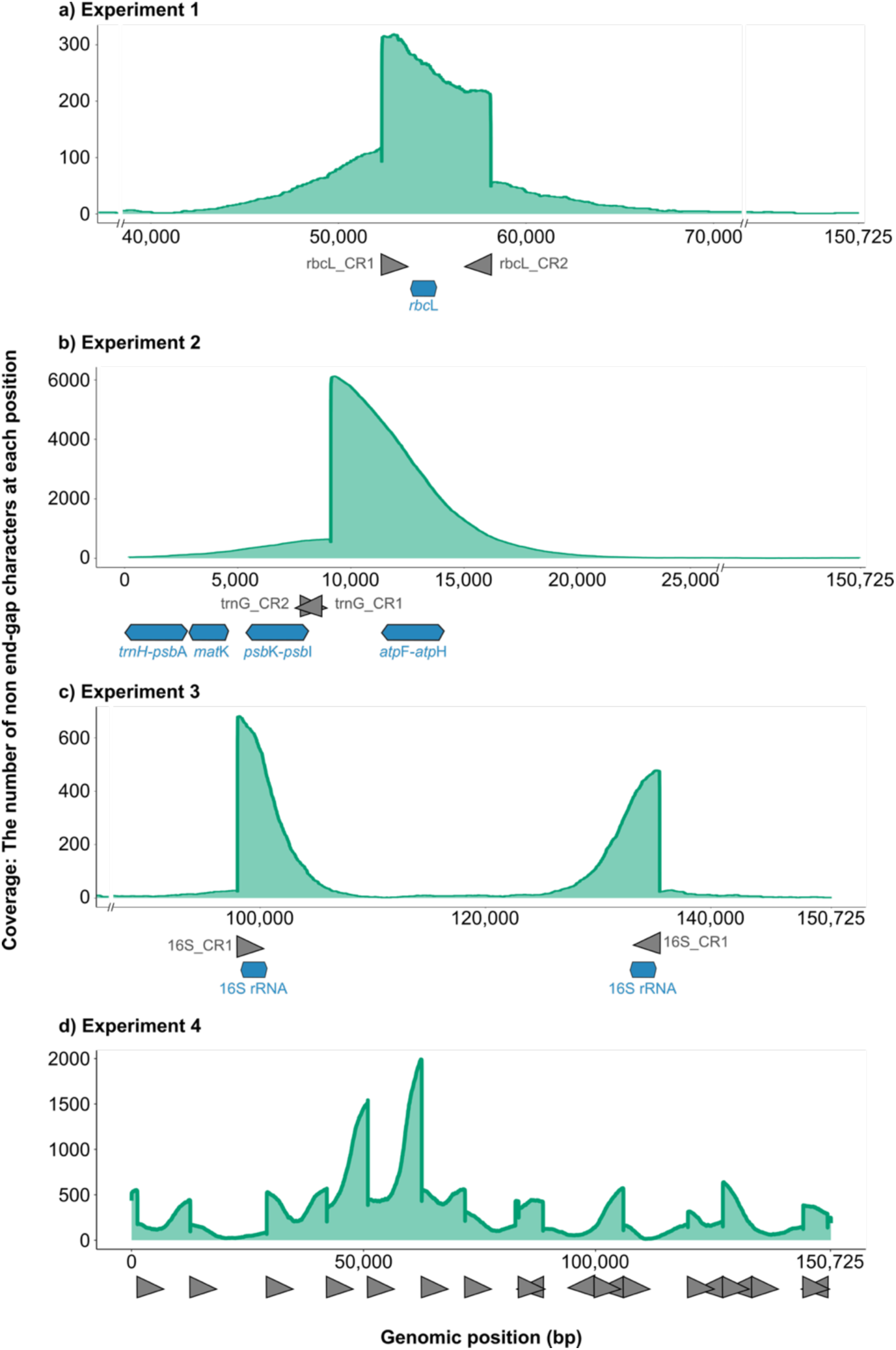
Coverage of raw sequence reads that passed Guppy basecalling (Q ≥8) and were mapped to the spinach reference genome. In experiments 1-4, we enriched for **a)** the *rbc*L plant barcode region, **b)** multiple standard plant barcodes including *mat*K and *trn*H-*psb*A, **c)** the 16S rRNA inverted repeat regions, and **d)** the whole chloroplast genome. In panels **a)** to **c)**, blue hexagons indicate the positions of target barcodes in the spinach reference chloroplast genome. In all panels, gray arrowheads identify the gRNAs binding sites (Table 1). Gaps in non-target sections of the reference genome are shown using –//– notation.

**Figure 4.**
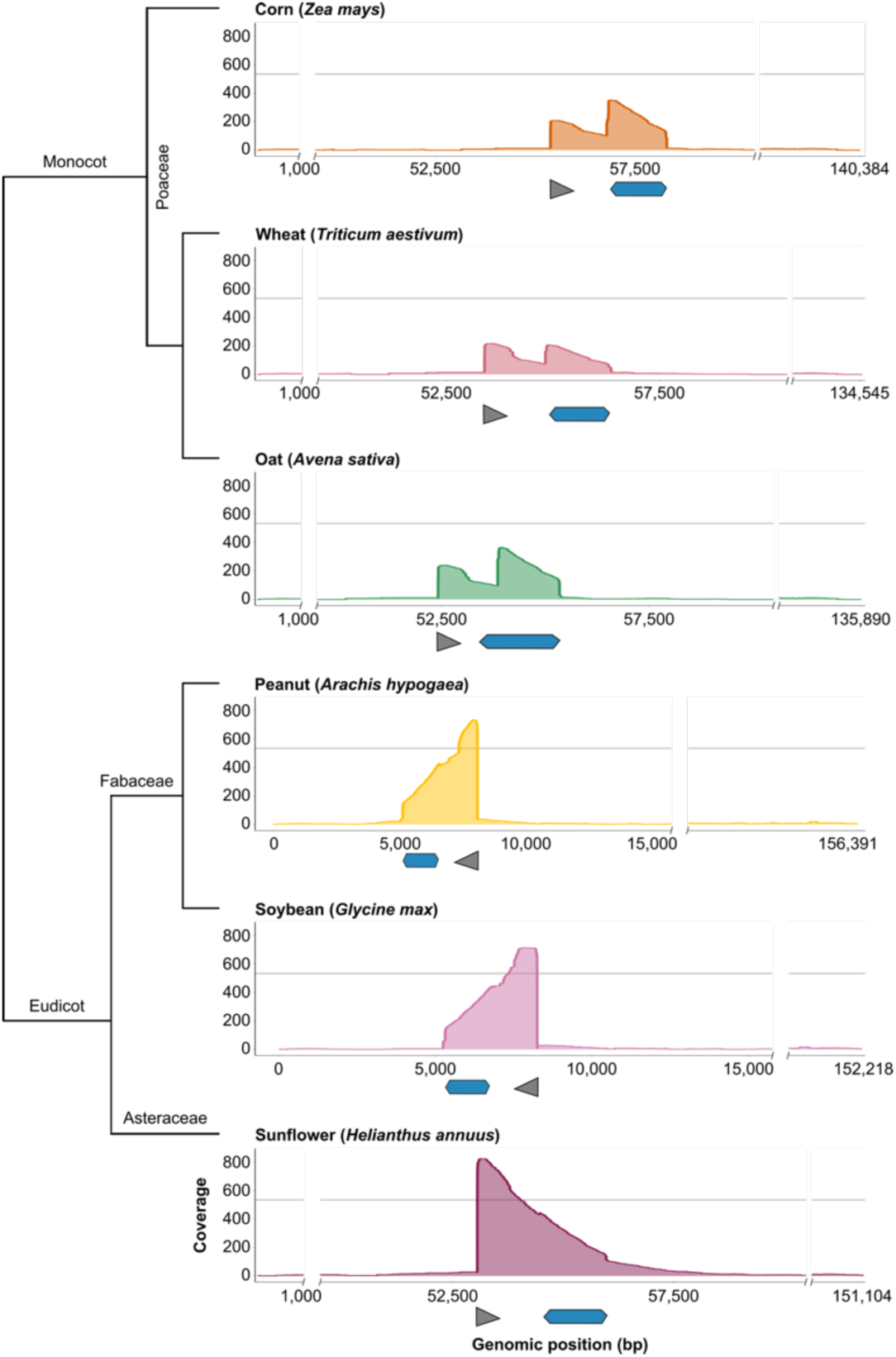
Depth of coverage for raw sequence reads that passed Guppy basecalling (Q ≥8) and mapped to reference genomes in mixed-species Experiment 6. The phylogeny of the six target taxa is shown to the left. Blue hexagons indicate the position of the *rbc*L gene, gray arrowheads indicate where the gRNA binds in each target-taxa, and the coverage values represent the number of non-end-gap characters obtained from sequences mapping to each position. The gray horizontal lines indicate the expected depth of sequence coverage that would have been obtained for each target taxon under the assumption that the 6 target species have equal DNA relative abundances. Gaps in the off-target portions of the reference genomes are indicated using –//– notation. Peak coverage differs in location in each target-taxon due to different chromosomal arrangements across taxa. The gRNA used (*rbc*L_CR1) was predicted to bind ∼1,500 bp up- or downstream from *rbc*L in each reference genome.

Experiment 2 targeted multiple plant DNA barcodes with overlapping forward and reverse gRNAs, yielding a total of 10,506 reads with a Q-score ≥8 (Table 2, Figures 2-3). Raw sequence lengths ranged from 126 bp to 19,890 bp (mean: 4,149 bp); when mapped to the reference spinach chloroplast, most raw reads sat downstream of the forward gRNA (Figure 3b) which was unexpected given that two gRNAs were used that ran in opposite directions from a single enrichment site. In total, 38 error-corrected reads were produced; 33 (87%) of these corrected consensus reads mapped to the spinach chloroplast reference genome. Of the corrected consensus reads that mapped to the spinach chloroplast reference genome, 32 reads aligned downstream of the forward gRNA and 1 read overlapped the forward and reverse gRNA (Table 2). *De novo* assembly generated 1 contig of 10,062 bp that aligned to the target region in the spinach chloroplast reference genome at 32X coverage and 99.2% identity to the reference (Table 2). When mapped to the spinach chloroplast reference, the assembled contig sat downstream of the forward gRNA and therefore only included one (*atp*F-*atp*H) of the four target barcodes (*mat*K, *trn*H-*psb*A, and *psb*K-*psb*I not included).

Experiment 3 targeted the 16S rRNA inverted repeat region using a single gRNA. A total of 3,117 reads had a Q-score ≥8 with a mean sequence length of 4,023 bp (range: 139-22,583 bp) which span the length of the 16S rRNA region (Table 2, Figures 2-3). A total of 54 error-corrected reads were produced; 34 (63%) of these corrected consensus reads mapped to the spinach chloroplast reference and all aligned to the target region (Table 2). *De novo* assembly generated 1 contig of 8,911 bp that mapped to the correct two locations within the chloroplast genome with 31X coverage and 99.4% identity to the reference sequence (Table 2).

### Whole chloroplast approach

The CRISPR-based enrichment approach yielded high sequencing depth of coverage and accuracy in sequencing the spinach chloroplast genome (Experiment 4). We obtained 15,766 reads with a Q-score ≥8 and a mean read length of 4,328 bp (range: 109-25,372 bp; Table 2, Figures 2-3). A total of 1,256 error-corrected reads were produced and 1,118 (89%) of these mapped to the spinach chloroplast reference genome (Table 2). *De novo* assembly generated 9 contigs of 1,733-18,510 bp that all aligned to the reference genome (Table 2). Of the 9 contigs, 2 contigs occurred in the inverted repeat regions. Together, the 9 contigs covered 81% of the spinach chloroplast reference genome (121,284 bp of 150,725 bp) with an average 60X coverage (20X – 128X coverage across contigs) and provided excellent accuracy with 99.3-99.6% identity to the reference genome (Table 2).

### Mixed-species approach

*In vitro*, we had varied depths of sequence coverage and accuracy in sequencing *rbc*L from a mixed set of 6 target species: soy, wheat, corn, peanut, sunflower and oats (Experiments 5-6). For Experiment 5, two gRNAs were used, a total of 7,569 reads had a Q-score ≥8 (Table 2, Figure 2) with a mean sequence length of 1,853 bp (range: 101-13,064 bp), and 22 error-corrected reads were produced (Table 2). Between 14 and 19 of these consensus-corrected reads mapped to each target-taxon’s chloroplast reference genome, and 2 *de novo* contigs were generated using metaFlye (2,827 bp, 6X coverage; Table 2). These 2 contigs mapped to the six reference genomes with pairwise identities of 80.8%-86.9%, but with only two contigs we inevitably failed to recover the full taxonomic breadth of six species included in the samples (Table 2). We therefore tried a second approach to contig assembly where trimmed reads were corrected and assembled into contigs independently for each target taxon and using this taxon-dependent approach we obtained 63-102 consensus (error) corrected reads for each taxon and we were able to build contigs that that were often more accurate with 79.2-98.6% identity to the reference chloroplast genomes at 25X-37X coverage (Table 2). Within the standard *rbc*L barcode region (553 bp), we found better accuracy (86.9-99.5%) between the contigs and the target chloroplast genomes.

In Experiment 6, we used only a single gRNA to sequence *rbc*L from the mixed sample and we obtained 39,507 reads that had a Q-score ≥8, which was a 5.2-fold larger value than the prior experiment that used two gRNAs (Table 2, Figures 2, 4). Mean sequence length was 1,627 bp (range: 130-18,342 bp). Similar to the prior experiment, we found that building contigs independently for each target-taxon resulted in better pairwise identity and taxonomic breadth (Table 2). For each target-taxon, between 115-179 error-corrected reads were produced and between 1-3 contigs were built which mapped to the chloroplast reference genome of each species with 73.2-99.4% identity and 28X-66X coverage (Table 2). Similar to Experiment 5, the portion of these contigs that spanned the standard *rbc*L barcode yielded better accuracy (86.7-100%) compared to the reference genomes. Thus, when comparing our methods based on 1 vs. 2 gRNAs to target a region of interest in a mixed-species sample, we obtained better depth of coverage with 1 gRNA but better accuracy with 2 gRNAs (Table 2).

Our final goal for the mixed-species approach was to compare CRISPR- and PCR-based methods for estimating DNA sequence relative read abundance. Both Experiment 5 (2x gRNA approach) and Experiment 6 (1x gRNA approach) produced contigs corresponding to all 6 taxa in relatively even proportions compared to PCR (Figure 5). The CRISPR-based strategies yielded strikingly accurate estimates of DNA relative read abundances for the three taxa of equal (1/6^th^) biomass proportions (Experiment 5: oat, 19%; sunflower, 15%; peanut, 13%; Experiment 6: oat, 22%; sunflower, 19%; peanut, 12%) compared to PCR (oat, 1%; sunflower, 35%; peanut, 38%; Figure 5).

**Figure 5.**
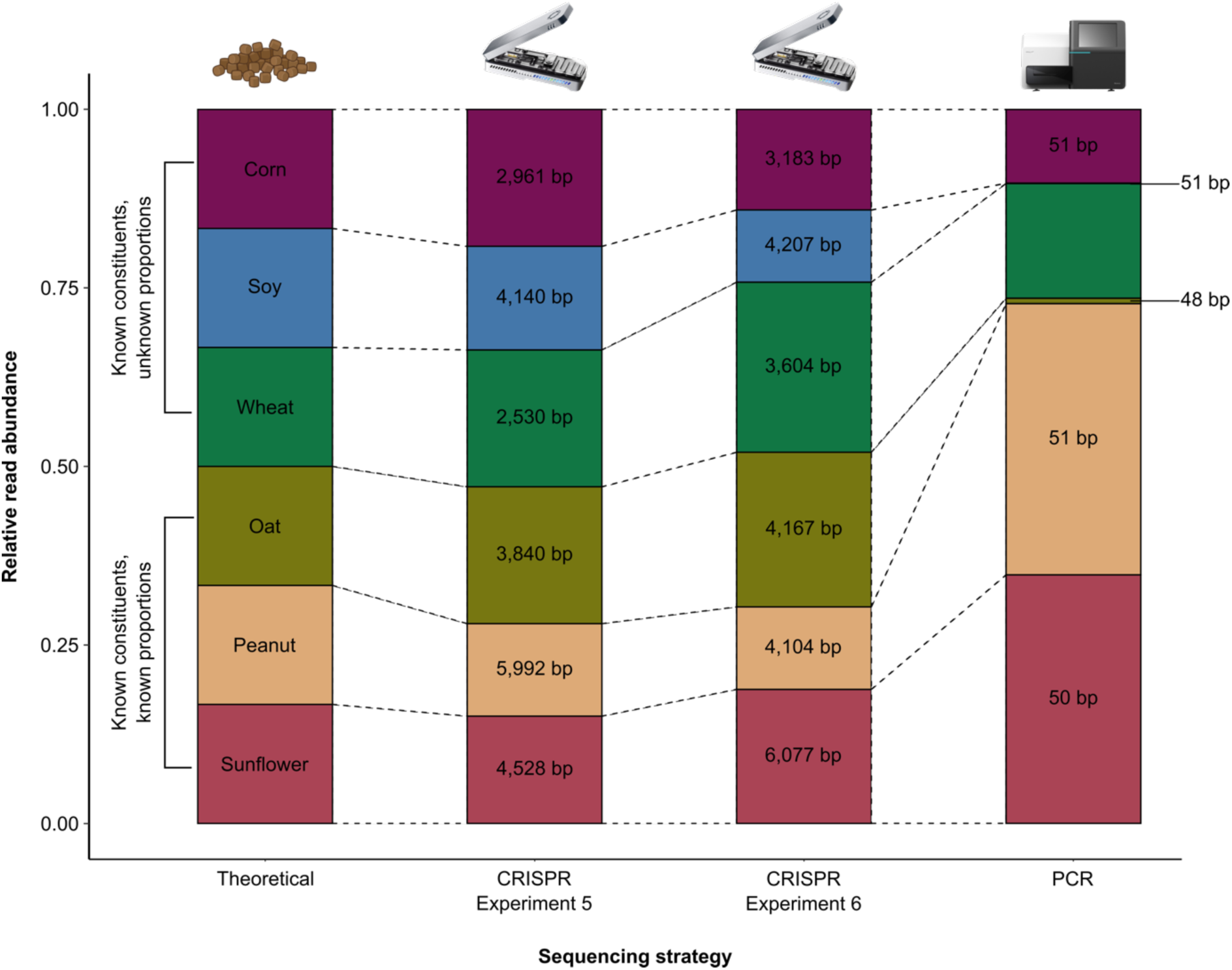
Stacked barplots comparing the results of CRIPSR- and PCR-based methods. From left to right, we show: the hypothetical relative biomass of each target-taxa in the mixed-species sample, contig coverage for Experiment 5 of CRISPR-Cas enrichment (2 x gRNA), contig coverage for Experiment 6 of CRISPR-Cas enrichment (1 x gRNA), and relative read abundance obtained using PCR-based DNA metabarcoding. Of 6 taxa in the mixed sample, 3 were of known proportions (oat, peanut, and sunflower; ⅙ each) and 3 were of unknown proportions (corn, soy, wheat). All target taxa were detected with CRISPR-Cas enrichment, which produced mean contig lengths per taxon that were at least 66-fold longer than amplicon sequencing.

## Discussion

Although CRISPR is generally underutilized in environmental science (Phelps et al., 2020), CRISPR-based enrichment strategies have shown promise and versatility (Baerwald et al., 2023; López-Girona et al., 2020; Ramon-Laca et al., 2022; Sánchez et al., 2022; Sandoval-Quintana et al., 2023; Williams et al., 2023). We evaluated strategies to harness this power for important applications such as overcoming the plant DNA barcode resolution problem (CBOL Plant Working Group, 2009; Kress, 2017) and issues with PCR-based DNA relative read abundance calculations (Deagle et al., 2019; Littleford-Colquhoun, Freeman, et al., 2022). Here, we were able to (*i*) identify many broad-spectrum gRNAs within the chloroplast genomes phylogenetically disparate and economically important taxa, (*ii*) demonstrate methodological versatility for sequencing plant DNA barcode loci, (*iii*) enrich and assemble a nearly complete chloroplast genome using just 12 gRNAs, and (*iv*) profile plant DNA within a mixed sample with remarkable accuracy and precision. Success with some of these CRISPR-Cas applications were relatively straightforward, but others, while promising in many regards, revealed opportunities for improvement and general points of caution for future work.

This study demonstrated the versatility of CRISPR-based enrichment approaches that extend beyond taxon-specific detection to include ‘universal’ methods that work across a broad swath of the plant phylogeny. Initially, we identified a total of 54,837 gRNAs across all 7 target taxa (Supplementary Data Table 1). Of these candidate gRNAs, 44,510 occurred in only 1 of the 7 target plant taxa with 100% identity (Supplementary Data Table 1) and provided one or more enrichment sites, which could potentially enable taxon-specific detection and sequencing of economically and ecologically relevant species. Additionally, 10,327 of the candidate gRNAs were identified in ≥2 target species and could provide one or more enrichment sites across multiple phylogenetically diverse species, including 398 broad-spectrum gRNAs that were identified in all 7 target species (Supplementary Data Table 1). In addition to validating several methods to enrich widely used plant DNA barcodes from a sample of a single species (Experiments 1-3), we also succeeded in multiplexing gRNAs to sequence most of the spinach chloroplast genome (Experiment 4). While whole genome assembly using CRISPR enrichment may have advantages, we need to assess this directly (Johnson et al., 2019). Perhaps most promisingly, we demonstrated the feasibility of sequencing mixed-species samples (Experiments 5-6) and showed that the CRISPR-based strategy to sequence the *rbc*L barcode from a mixed sample delivered both greater lengths and more accurate relative abundances than a typical PCR-based method. It is widely acknowledged that PCR-based methods can result in stochastic and biased abundance data (Pawluczyk et al., 2015), with strategies used to process such errors remaining largely contentious within the field (Littleford-Colquhoun, Freeman, et al., 2022; Littleford-Colquhoun, Sackett, et al., 2022); therefore, if CRISPR-Cas enrichment is capable of producing more accurate relative read abundance data, and we are able to collectively address opportunities for improvement, much of the consternation within the field could be eliminated using this alternative strategy. Comparison of our two mixed-species enrichment experiments raises the question of whether methods that utilize one gRNA vs. a multiplex of two or more gRNAs generally present the type of tradeoff in the potential the depth of sequence coverage vs. sequence accuracy that we observed needs to be explored in other systems. Establishing general expectations about the efficacy of different mixed-species approaches could be beneficial to applications that require especially precise taxonomic determinations or relative abundance estimates from environmental DNA.

While it is relatively straightforward to develop gRNAs for single-species enrichment and sequencing (Knott & Doudna, 2018), translating these protocols to mixed-species applications for use with environmental DNA will require careful planning and analysis. While we encountered off-target enrichment in all experiments, suggesting the gRNAs had some functionality at other loci (e.g., the nuclear genome), this off-target activity was especially problematic in the mixed-species data generated for samples that contained more highly processed, potentially degraded, and genetically diverse template DNA (Experiments 5-6). With these mixed-species experiments, a lower percentage of reads that passed the Guppy basecaller mapped back to any of the reference genomes for species that were included in the mixed-species sample (Figure 2). Further complicating matters, only a fraction of the total reads from each run map to reference genomes in mixed-species analyses—this reduced per-species coverage presents a challenge that will be amplified in more complex samples that include a greater diversity of both target and non-target DNA. For instance, while many previous studies have shown some degree of off-target enrichment when using single-species samples (López-Girona et al., 2020; Ramon-Laca et al., 2022), Sandoval-Quintana et al. (2023) found that only 0.03% of good quality reads covered the region they wished to study when enriching a bacterial gene from a complex microbial sample. To address this challenge, it may be useful to experiment with mismatch tolerance values. For example, by allowing ≤2 bp mismatch tolerance at the 5’ end of the gRNAs in our analysis, we were able to expand the number of candidate gRNAs for testing across the seven species used in these trials. But there will be a tradeoff between mismatch tolerance and off-target activity that needs to be considered for the types of mixed-species samples that are often used for analyses of environmental DNA.

Perhaps the greatest need for further research required to translate CRISPR-based enrichment methods for the sequencing of complex mixtures will revolve around the strategies available to obtain on-target data and build contigs. Due to relatively low depth of coverage and percentage of base-called reads that mapped back to the reference genomes in Experiments 5-6, we were unable to construct species-specific contigs using metaFlye on the full dataset; we had to build contigs independently for each species using the raw reads that mapped back to each plant species’ reference genome. In many real-world applications involving environmental DNA, there will not be *a priori* knowledge of reference genomes for all species in the mixture (Yang et al., 2021) and thus more sensitive taxon-calling methods will be required. A promising strategy involves translating bioinformatic methods that are being developed specifically for the analysis of bacterial metagenome-assembled genomes (‘MAGs’) for future applications involving CRISPR-based enrichment sequencing (Parks et al., 2017; Stewart et al., 2019; Tully et al., 2018), but our results suggest that achieving acceptable levels of accuracy may ultimately require more than simply ‘tuning’ the parameters used in these existing methods (e.g., Experiments 5-6). Similarly, computational methods that facilitate upstream screening gRNAs for off-target activity across multiple taxa could improve options available to overcome these downstream challenges. To our knowledge, no such cross-taxon site selection tool is currently available.

Considerations for future experimental design at the bench could help enhance the versatility and accuracy of CRISPR-based sequencing methods for biodiversity research, especially for challenging mixed-species analyses. For example, DNA extraction methods (Kang et al., 2023; Russo et al., 2022), the number of purification steps used during library preparation (De La Cerda et al., 2023), the specific Cas system deployed (e.g., Cas9 vs. Cas3 or Cas12a; Schultzhaus et al., 2021), and the sequencing platform utilized (e.g., Oxford Nanopore vs. Illumina or PacBio; Li & Harkess, 2018) may need to be optimized in order to ensure adequate on-target sequence coverage. Encouraging strategies to enhance enrichment of on-target reads include methods for tiling gRNAs, whereby overlapping gRNAs can be used to extend the enrichment of the target region (López-Girona et al., 2020), or to improve the median depth of coverage for a particular locus (Gilpatrick et al., 2020). In addition, there are encouraging strategies to deplete non-target sequences, such as host DNA in microbiome studies, using CRISPR-Cas selective amplicon sequencing (Zhong et al., 2021).

Our *in-silico* analysis of plant gRNAs coupled with our six validation experiments provide a proof of concept involving the use of CRISPR-based enrichment sequencing for use in environmental biology. This technology can be used to build accurate plant DNA barcode libraries with sequences that are long enough to span multiple barcode regions and thus overcome long-standing limitations to taxonomic resolution in PCR-based barcoding studies (CBOL Plant Working Group, 2009; Kress, 2017)—potentially providing sequences for entire chloroplast genomes—though overcoming the challenge of translating this potential into versatile and cost-effective methods for analysis of environmental DNA represents an exciting area for development (Ramon-Laca et al., 2022; Schultzhaus et al., 2021). Moving forward, these approaches can be extended to incorporate other facets of research that are integral for biodiversity discovery, such as determining structural rearrangements (Sun et al., 2022), phylogenetic patterns (Yang et al., 2014), targeted genome sequencing (López-Girona et al., 2020), multiplexing samples and loci within a single reaction, and building metagenome-assembled genomes (Liu et al., 2022; Sandoval-Quintana et al., 2023).

## Acknowledgements

We would like to thank Colin Donihue, Alex Harkess, and Timothy Divoll. Funding was provided by IBES seed funding at Brown University and NSF DEB-2046797.

## Data availability

Nanopore sequence read data and sample metadata for all experiments have been made available at NCBI (BioProject: PRJNA989586). Bioinformatic steps taken for each experiment can be found on the Brown Digital Repository (https://repository.library.brown.edu/studio/item/bdr:482c9vvy/). Illumina sequence read data and sample metadata for the mixed-species sample have been made available at NCBI (BioProject: PRJNA989255). Files used for building the global plant reference library can be found on the Brown Digital Repository (DOI to be provided upon manuscript acceptance).

## Notes

https://repository.library.brown.edu/studio/item/bdr:482c9vvy/

### Competing Interest Statement

The authors have declared no competing interest.

