## Appendix 1 for "A CRISPR-based strategy for targeted sequencing in biodiversity science"

**Manuscript category:** Resource Article

**Short title**: Versatile CRISPR-based biomarkers

**Authors**: Bethan Littleford-Colquhoun[1,2,*] & Tyler R. Kartzinel[1,2,*]

**Affiliations**:

[1] Department of Ecology, Evolution, and Organismal Biology, Brown University, Providence, RI 02912, USA

[2] Institute at Brown for Environment and Society, Brown University, Providence, RI 02912, USA

**Appendix S1**

*PCR and Illumina sequencing of the mixed-species sample*

To compare the output of the CRISPR-Cas9 approach to metabarcoding to PCR, we used Illumina sequence data generated for BioProject PRJNA989255 which included the sample used for the mixed-species CRISPR-Cas approach (GFT105.2) and its corresponding control samples (extraction blank, PCR positive and negative control).

The Illumina data for this BioProject was generated by amplifying the well-known *trn*L-P6 marker for plant DNA in 25 µL reactions comprising 1.5 mM $MgCl_2$, 200 µM each dNTP, 0.2 µM each primer [*trn*L(UAA)g / *trn*L(UAA)h; (Taberlet et al., 2007)], 1X Platinum Taq buffer (-$MgCl_2$), Platinum Taq DNA polymerase (0.5 U/rxn), molecular water, and 2 µL of DNA extract. The *trn*L-P6 marker is often used for plant metabarcoding studies due to its shorter length, conserved primer sites, and interspecific variation (Taberlet et al., 2007). The primers were modified to contain Illumina Nextera Transposase Adapters overhangs. Thermocycling followed a program of initial denaturation at 95°C for 5 min, followed by 35 cycles of 95°C for 30 s, 55°C for 30 s, and 72°C for 1 min, with a 10 min final extension at 72°C. Amplicons were visualized by running 2 µl of PCR product on a 1.5% agarose gel and PCR success was verified by confirming the presence of a band for the samples and the PCR positive controls (*Spinacia oleracea*) and the absence of a band for the extraction blanks and PCR negative controls (molecular water). To obtain DNA sequences, amplicons were multiplexed using the Illumina P5/P7 adapters in a second round of PCR prior to sequencing at the RI-INBRE Molecular Informatics Core. Samples were pooled at equimolar concentrations and sequenced on the Illumina MiSeq platform with 2 x 150 bp paired-end chemistry. The resulting sequences were demultiplexed and adapter sequences were removed for analysis via BaseSpace using standard Illumina FASTQ generation pipelines.

In order to identify the DNA sequences, we constructed a global reference library using *trn*L-P6 sequences available from the European Nucleotide Archive (ENA). We obtained all plant sequences from the ENA 143 release (N = 599,992 accessions) and extracted full-length *trn*L-P6 sequences using the *ecoPCR* function by searching for sequences within the *g* and *h* primers that were 8-300 bp and allowing a maximum of 3 mismatches to each primer. This global reference library included 21,422 unique *trn*L-P6 sequences from 156,880 entries and represented at least 615 plant families.

For the demultiplexed Illumina sequence data generated for BioProject PRJNA989255, we removed primers from forward and reverse reads using *cutadapt*, allowing maximum of 3 mismatches and no indels (Martin, 2011). Only paired-end sequence reads where both primer sequences could be identified in both directions were assembled using the *illuminapairedend* in obitools with a minimum alignment score of 40 and only overlapping assembled sequences retained for further analysis (Boyer et al., 2016). To ensure quality control of sequence data, we ensured that positive controls and samples yielded relatively large numbers of sequences compared to extraction blanks and negative controls. We used the *obiuniq* command to tally and merge identical sequences. Sequences that occurred <2 times overall or that were outside of the broad range of expected sequence lengths for the marker (<8 bp or >300 bp) were discarded using the *obigrep* command. Sequences were considered to be likely PCR artifacts if they were highly similar to another sequence (1 bp difference) and had a much lower abundance (0.05%); we discarded these sequences using the *obiclean* command. We identified each of the unique sequences in the dataset using the global reference library based on the *ecoTag* command in obitools. This analysis generated a file representing a table of sequence counts per food-taxon per sample and a table containing the taxonomy of each food-taxon from which we then extracted the mixed-species sample (GFT105.2) for analysis; a total of 85,940 high quality reads representing 337 unique sequences were retained in the mixed-species sample (Supplementary Data Table 2).

We took two approaches to analyzing the Illumina sequence data for the mixed-species sample. First, we eliminated sequences that did not perfectly match a reference sequence in the global library (100% identity; 33 sequences retained). Of these, some had ambiguous species-level identifications, requiring us to review individual BLAST results to determine if one of the target-taxa (corn, oat, wheat, soy, peanut, sunflower) was listed among top taxa or if it should be removed as a putative error (19 sequences retained). The read counts for sequences that were identified as being from the same target-taxon were then combined and resulted in a total of 82,483 reads representing the 6 target-taxa. Second, we relaxed the 100% matching requirement due to the possibility that not all haplotypes of the target-taxa present in the mixed sample were also represented in the global reference library. By reducing the threshold to 92% and classifying sequences only to the three major plant lineages included in the mixed-species sample (i.e., monocots, superrosids, and superasterids; Figure S1), we were able to evaluate the sensitivity of our relative read abundance estimates to uncertainties surrounding the taxonomic precision of short-read sequences.

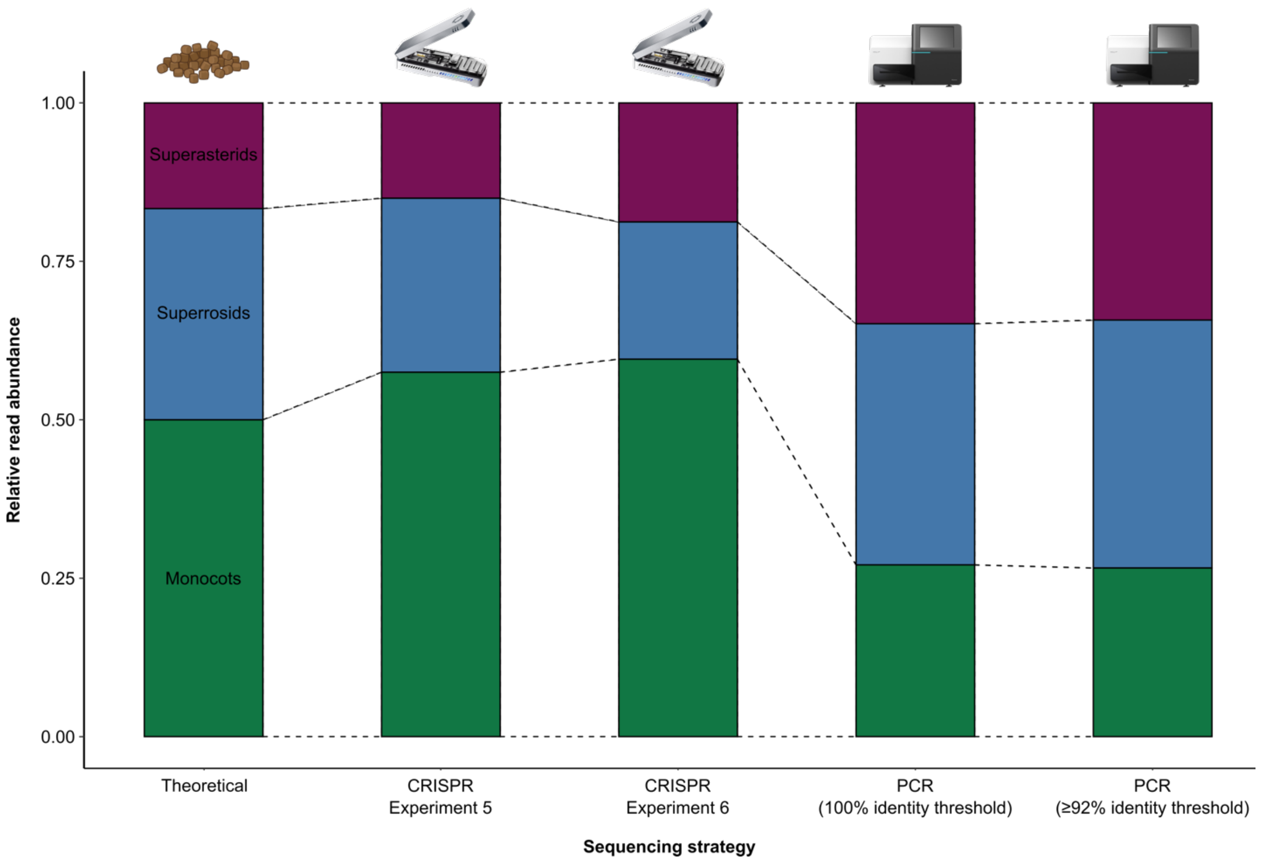


**Figure S1.** Stacked barplot of the hypothetical relative biomass of the three major plant groups present in the mixed-species sample, contig coverage for Experiment 5 of CRISPR-Cas enrichment, contig coverage for Experiment 6 of CRISPR-Cas enrichment, the relative read abundance obtained using a strict (100%) sequence identity threshold for PCR-based method for DNA metabarcoding, and the relative read abundance obtained using a relaxed (≥92%) sequence identity threshold for PCR-based method for DNA metabarcoding. Both CRISPR-based experiments produced contig coverage in more-similar proportions to theoretical proportions compared to PCR-based methods. When comparing PCR-based methods, the relative read abundance for the three major plant groups present in the mixed-species sample were similar regardless of identity threshold used (100% vs. ≥92%).
